# Bioactivity Analysis of Secondary Metabolites from Actinomycetes’ Isolated from Coastal Sediments

**DOI:** 10.1101/2024.09.08.611773

**Authors:** Kumar Shubhro, Arifa Afrose Rimi, Md. Emdadul Islam, Kazi Didarul Islam, Md. Morsaline Billah, Mijan Mia, SM Mahbubur Rahman

## Abstract

The rising threat of antibiotic resistance underscores the need for new antibiotics. Marine actinomycetes have emerged as promising sources of bioactive compounds. In this study, 22 isolates from marine sediments were morphologically identified, with nine confirmed as actinomycetes through 16S rRNA gene sequencing. Five strains- *Streptomyces olivoverticillatus* (T-2), *Streptomyces cyaneus* (T-4), *Nocardiopsis synnemataformans* (T-7), *Streptomyces albogriseolus* (T-8), and *Streptomyces atrovirens* (T-9)- exhibited significant antibacterial activity. Cultured in Starch Casein Broth, their metabolites were tested for antibacterial, antioxidant, anticoagulation, and anti-inflammatory activities. T-4 and T-8 demonstrated notable antimicrobial action, T-8 showing potent DPPH radical scavenging (372.09 ± 11.05 µg/mL). T-9 inhibited trypsin (IC_50_ 435.12 ± 15.88 µg/mL) and had a prothrombin time of 12.08 ± 1.46 minutes. T-8 enhanced RBC membrane stabilization (IC_50_ 140.08 ± 2.30 µg/mL). These findings suggest that marine sediment-derived actinomycetes hold significant therapeutic potential and warrant further study.

## Introduction

Actinomycetes are a group of gram-positive, aerobic, spore-forming bacteria that belong to the Actinomycetales order and resemble fungi (Arifuzzaman et al., 2010). They are widespread not only inside the boundaries of the territory but also in aqueous environments such as freshwater and the ocean (Valli et al., 2012). This group of bacteria is mainly recognized as the potential producer of essential bioactive compounds like antibiotics, antifungals, antioxidants, etc (Sengupta et al., 2015). More than 140 species of actinomycetes have been reported to have produced over 7,000 bioactive chemicals, with *Streptomyces spp*. They alone account for almost 80% of these compounds (Jensen et al., 2015). About 500 species of *Streptomyces* produce nearly two-thirds of currently available antibiotics (Mohanraj & Sekar, 2013) Peela et al. (2005). Some Actinomycetes that produce antibiotics have been isolated from marine environments; most of these isolates are *Streptomyces species*.

Antibiotic resistance arises from multi-drug-resistant bacteria responsible for numerous infectious, deadly diseases. The incidence of these infections is rising rapidly and at a concerning pace (Cassell & Mekalanos, 2001). Multiple drug resistance in various pathogenic bacteria is becoming difficult or sometimes untreatable with commercially available antibiotics (Frieri et al., 2017). While antibiotic resistance continues to increase, the development of new antibiotics is advancing slowly enough to meet the demand. Over time, even new antibiotics can become ineffective against bacterial infections. For instance, multi-drug resistance in carbapenemase-producing Gram-negative bacteria, gonorrhea, and tuberculosis is a significant concern in modern medical science. That is why antibiotic resistance becomes a major threat to human beings (MacGowan & Macnaughton, 2017). So, a safe and effective new antibiotic is required to fight against antibiotic-resistant bacteria (Newman & Cragg, 2016). Marine microorganisms like actinomycetes show enormous chemical diversity, which can be used to treat this kind of disease.

The marine environment has proved an effective ecosystem to produce essential novel secondary compounds (Attimarad et al., 2012). Over 70% of the Earth’s surface is ocean, and marine ecosystems are more diverse than land (Attimarad et al., 2012). Since actinomycetes are known producers of bioactive compounds, new and novel compounds can likely be discovered (Hughes & Fenical, 2010). This research aims to separate actinomycetes that produce antibiotics from coastal sediments in Bangladesh’s Vola District and evaluate their antibacterial qualities.

## Materials and Methods

### Collecting soil samples

Soil samples were gathered from multiple locations within Bangladesh’s Char Kukri Mukri and Char Montaz districts. Using a sterile spoon obtained samples from a 15-20 cm depth in the coastal sediments. After that, the samples were put into sterile falcon tubes and immediately sealed in an ice box. Pure cultures of actinomycetes were obtained (Poosarla, 2013).

The samples were heated for two hours at 60°C after being dried to remove any undesirable microbes. After that, a test tube containing one gram of each soil sample and ten milliliters of sterile distilled water was shaken for approximately five minutes. Then, serial dilutions were made between 10^-1^ and 10^-6^. The diluted samples were added to Petri dishes with Starch-Casein Agar medium treated with vancomycin to decrease bacterial growth and cycloheximide to prevent fungus development. For five to seven days, the plates were then incubated at 30°C.

### Isolation of genomic DNA

Actinomycetes’ genomic DNA was obtained using a slightly modified version of the (A et al., 2018) procedure. Actinomycete strains were grown in nutritional broth at 30°C in shaking incubators for three days. Centrifuging at 13,000 rpm for 5 minutes after incubation produced the bacterial pellet containing 15-20 mg of cells. 50 µL of 10% SDS, 100 µL of 20 mg/mL lysozyme, ten mM EDTA, 2.5 M NaCl, 3.5% CTAB, and 0.1 M Tris-HCl pH eight were used to lyse the pellet. Once 20 µL of 20 mg/mL RNase was added, the mixture was vortexed for 5–10 minutes and then incubated for 30 minutes at 37°C. Each proteinase-K addition was followed by a 30-minute incubation period at 65°C.

The mixture was centrifuged, and the supernatant was transferred into a fresh tube. After adding a 24:1 chloroform-isoamyl alcohol mixture and transferring the aqueous phase into a fresh Eppendorf tube containing 0.8 volumes of isopropanol, another centrifugation was carried out.

Two hours at -20°C incubation were followed by centrifugation, 70% ethanol washing, air drying, and resuspension of the DNA pellet in 50 µL of TE buffer. The DNA’s concentration and purity were assessed using a Nanodrop spectrophotometer, and the results were verified by 1% agarose gel electrophoresis. After that, the DNA was stored at -20°C.

### 16s rRNA gene amplification

Using primers C27 (5’-AGAGTTTGATGGCTCAG-3’) and RC1492 (5’-TACGGCTACCTTGTTACGACTT-3’), obtained from Gene DireX, the actinomycetes’ 16S rRNA gene was amplified. Supplement File Table 1 provides information on the chemicals and concentrations used in the 50 µL Polymerase Chain Reaction (PCR) combination. At the same time, Supplement File Table 2 details the PCR cycle parameters and exposure periods.

### Identification of the bacteria

The PCR products were sequenced once they were obtained. The resulting sequences were evaluated using a BLAST search at the National Center for Biotechnology Information (NCBI) and then uploaded to GenBank. A phylogenetic tree was constructed using the neighbor-joining technique and MEGA X for multiple sequence alignment.

### Fermentation

According to Song et al. (2012), fermentation of the actinomycetes was done. After being infected with a loopful of actinomycetes colony, a 250 mL conical flask containing 100 mL of seed culture (made up of 10 g/L of starch, four g/L of yeast extract, and two g/L of peptone, pH 7.0) was cultured for 48 hours at 28°C. After that, the seed culture was moved to a 2 L flask containing 900 mL of fermentation medium (which included Starch 10 g/L, Casein 0.3 g/L, KNO_3_ 2 g/L, K_2_HPO_4_ 2 g/L, NaCl 2 g/L, MgSO_4_ 2 g/L, CaCO_3_ 0.02 g/L, and FeSO_4_.7H2O 0.01 g/L). The flask was then shaken, and at least four weeks of fermentation was allowed.

### Extraction of secondary metabolites

With some modifications, the secondary metabolites were extracted by Nurunnabi et al., 2018 and Gebreyohannes et al., 2013. After centrifuging the bacterial cells to extract the supernatant, ethyl acetate was added in a 1:3 ratio, and the mixture was allowed to rest for three days. The target extract-containing ethyl acetate layer was separated and kept dry using a funnel. The solvent was removed from the extract using a rotating vacuum evaporator and stored at -20°C.

### MIC determination

MIC determination was done according to Sarker et al. (2007). One pathogenic bacterial colony was cultivated in Mueller Hinton Broth (MHB), which contained 1.5 g/L starch, 2.0 g/L beef extract, and 17.5 g/L casein acid hydrolysate. After incubating for twenty-eight to twenty-four hours, the bacterial suspension was adjusted to 5 × 10LJ CFU/mL using the 0.5 McFarland standard. Standards and extracts were prepared at 1600 µg/mL.

Each well in a 96-well microtiter plate that had been sterilized received 100 µL of MHB. The test solution was divided among the first nine wells; the twelfth, tenth, and eleventh wells contained an antibiotic, a negative control, and a blank, respectively. About 18–24 hours were spent incubating at 37°C before adding 5 µL of resazurin dye. It was possible to determine the minimal inhibitory concentration (MIC) ten minutes later, thanks to the color shift.

#### Growth inhibition determination of the pathogens

The absorbance at 600 nm, indicating bacterial growth turbidity, was measured for each well. Next, using the given formula, the % inhibition rate was computed (Akinduti et al., 2019):

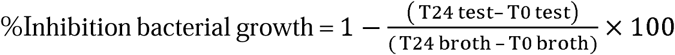

Where;

T24 test = Absorbance of the test sample (extract + test pathogen) at 24 hours

T0 test = Absorbance of the test sample (extract + test pathogen) at 0 hours

T24 broth = Absorbance of the broth culture (test pathogen + MHB) at 24 hours

T0 broth = Absorbance of the broth culture (test pathogen + MHB) at 0 hours

## Antioxidant Activity analysis

### The DPPH Free Radical Scavenging Assay

A slightly modified version of the Brand-Williams et al. (1995) method was used to measure the DPPH free radical scavenging activity. We calculated the % of inhibition using a formula attached below:

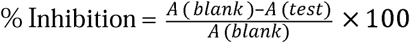

Where A (blank) is the absorbance of the blank and A (test) is the absorbance of the test solution

The IC_50_ figure indicates the concentration required to scavenge 50% DPPH free radicals. Plotting the % inhibition versus the sample extract concentration logarithm yielded this result.

### Reducing Power Assay

With some modifications, the reducing power was performed using Oyaizu’s method (1986). The following formula was used to get the % increase in reducing power:

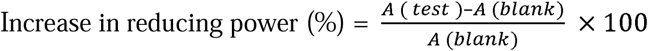

Where A(blank) represents the blank’s absorbance and A(test) the test solution’s absorbance, a graph showing the percentage inhibition against the logarithm of the sample extract concentration was used to get the IC_50_ value.

## Evaluation of Blood Coagulation Activity

### Serine Protease Inhibition Assay

Minor modifications were made to the Kunitz procedure (KUNITZ, 1946), and Serine Protease Inhibition was experimented. Two milliliter Eppendorf tubes were filled with 250 microliters of each of the five concentrations of the sample and standard (quercetin): 400, 200, 100, 50, and 25 µg/mL. Next, 500 µL of heated 1% casein solution and 250 µL of trypsin solution were added to each tube, and the tubes were incubated for 30 minutes at 37°C. After adding 300 µL of 5% trichloroacetic acid (TCA) to halt the reaction, the tubes were centrifuged for 15 minutes at 12,000 rpm. An absorbance measurement at 280 nm was used to calculate the percentage of inhibition.

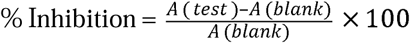

A(blank) indicates the absorbance of the blank, and A(test) indicates the absorbance of the test solution.

### Prothrombin Time Determination (PT) Assay

The procedure reported by Biswas et al., 2019 was used to determine the Prothrombin Time (PT) test. Ten healthy donors provided five milliliters of blood each, then placed into EDTA tubes. The tubes were centrifuged at 3000 rpm for fifteen minutes to extract the plasma. Subsequently, the plasma was transferred to fresh EDTA tubes and maintained at 4°C. 100 µL of the standard (Warfarin) at concentrations ranging from 50 to 800 µg/mL and 200 µL of plasma were mixed using a vortex mixer for each test. Upon adding 300 µL of 25 mM CaCl2, the tubes were incubated at 37°C. The coagulation durations were then monitored at 5-second intervals.

## Evaluation of the Anti-inflammatory Activity

### Human red blood cell (HRBC) membrane stabilization assay

The human red blood cell (HRBC) membrane stabilization assay was done according to Yesmin *et al*., 2020 with fewer adjustments. Five milliliters of blood drawn from healthy volunteers were placed in EDTA tubes and centrifuged for five minutes at 3000 rpm. After collecting the supernatant, the pellet was centrifuged for five minutes at 2500 rpm and washed three times with a sterile 0.9% NaCl solution. A 10% RBC solution was made using 154 mM sodium chloride and ten mM phosphate buffer (pH 7.4).

200, 100, 50, and 25 µg/mL of the sample and standard (Aspirin) were then serially diluted in phosphate buffer. In a 2 mL Eppendorf tube, 600 µL of isotonic solution, 600 µL of sample or standard, and 600 µL of RBC suspension were mixed. RBCs and a hypotonic solution were also prepared as controls. The samples were centrifuged for 15 minutes at 5000 rpm after a 10-minute incubation at 37°C. The absorbance at 540 nm was then measured. Next, the % of hemolysis inhibition was computed using the following:

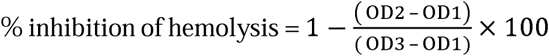

The test sample’s absorbance in an isotonic solution is measured as OD1, the control sample’s absorbance in a hypotonic solution is measured as OD2, and so on.

## Result

### Isolation and identification of Actinomycetes

Samples of soil were then serially diluted (10-² to 10LJLJ) and plated on Starch Casein Agar containing Vancomycin and Cycloheximide to isolate actinomycetes (see Supplementary Figure 1). Initially, 22 of the 47 bacterial isolates were picked, and ten were selected for additional examination based on their morphology (refer to Supplementary Figure 2). Starch Casein Agar cultivated and maintained these isolates as pure cultures. Genomic DNA was isolated for molecular identification, confirmed by 1% agarose gel electrophoresis (refer to Supplementary Figure 3), and quantified for purity and concentration using a NanoDrop spectrophotometer (Table 1). Figure 2 illustrates the PCR amplification of the 16S rRNA gene by using universal primers (C27 and RC1492). The accession numbers for the amplified sequences registered with GenBank are listed in Supplementary Table 3. Supplementary Table 4, BLAST findings and NCBI GenBank accession codes describe the molecular identification profiles. The isolated bacteria’s phylogenetic tree is shown in Figure 3.

**Figure 1:**
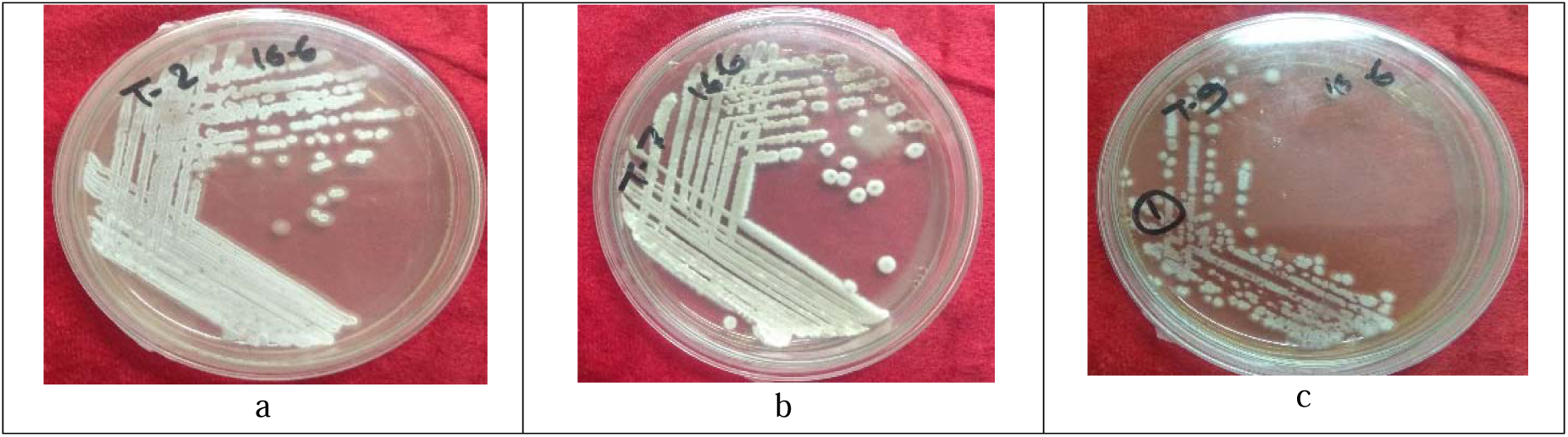
Pure culture of actinomycetes on SCA media cultured on petri dish a) *Streptomyces olivoverticillatus*, b) *Nocardiopsis synnemataformans*, c) *Streptomyces atrovirens*

**Figure 2:**
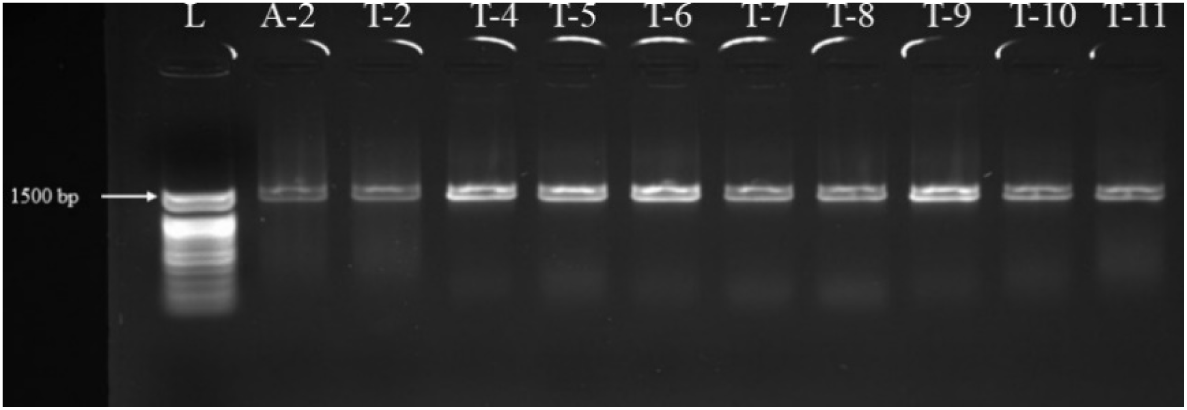
PCR amplification of the isolates’ 16s rRNA gene (5 µL PCR product was performed on 1% agarose gel)

**Figure 3:**
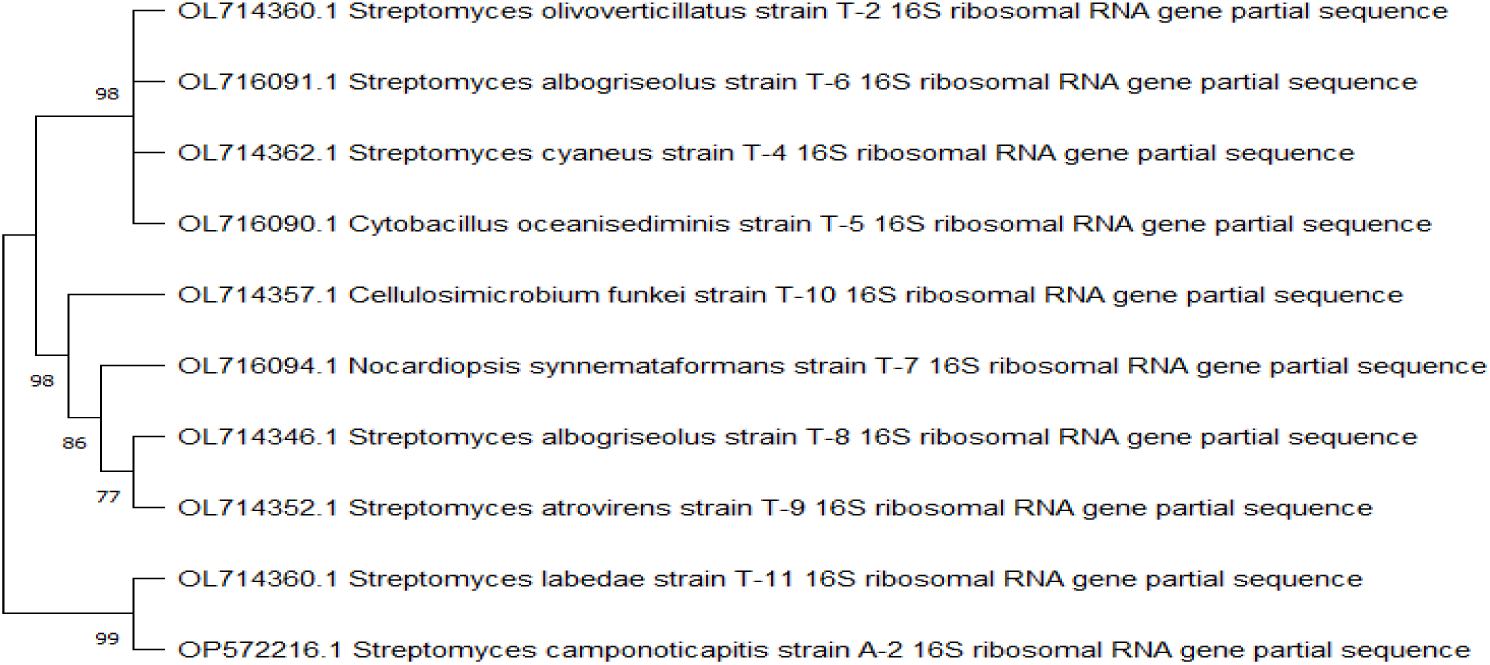
Using the neighbor-joining method in MEGA X software, a phylogenetic tree was built for the isolates from the Sundarbans based on 16S rRNA gene sequences.

### Antibacterial activity and MIC determination

Figure 4: How various extracts affect *S. typhi.* The first nine columns display extracts from several actinomycetes at concentrations ranging from 1600 µg/mL to 12.5 µg/mL. The negative control is the final column, the blank is the eleventh, and streptomycin is the positive control in the tenth column. There were three runs of this test. There is no need for statistical analysis because the Resazurin-based MIC test is qualitative. The maximum MIC value among the three tests is employed if there is a discrepancy of ± one-fold dilution in the results. Additional testing is necessary to validate inconsistent results.

**Figure 4:**
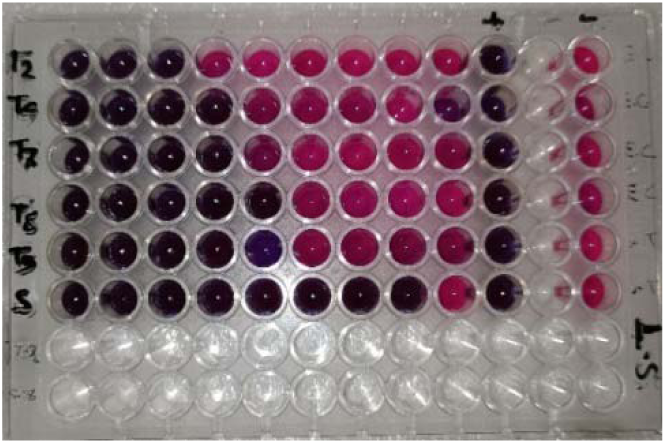
Resazurin-based MIC for antimicrobial activity against pathogenic bacteria.

This study has demonstrated that actinomycetes are among the most influential producers of antibacterial agents. They have been isolated from diverse soil samples, highlighting their role in the microbial diversity of ecosystems. Actinomycetes generate a range of biologically active compounds that perform various functions within their natural environments. (Janardhan et al., 2014). Coastal sediments exist in a distinctive environment with significant microbial and fungal competition. (Dilip et al., 2013). Our study agreed well with this previous study as 16s rRNA gene sequencing has helped identify them as actinomycetes. This study suggests that coastal actinomycetes can be potential microorganisms to help find novel antimicrobial compounds.

Medium polarity ethyl acetate dissolves polar and non-polar molecules to remove secondary metabolites from culture media efficiently. These chemicals are unaffected by reduced pressure rotary vacuum evaporation. Additionally, The antibacterial efficacy of five actinomycetes extracts against six pathogens is shown in Table 3. The minimum inhibitory concentration (MIC) was determined using resazurin, which remains purple in the absence of bacteria and turns pink when viable bacterial cells are present. Because of this characteristic, resazurin serves as a helpful gauge for determining how well the extracts stop the growth of bacteria (Karuppusamy & Rajasekaran, 2009).

The crude extracts obtained from the isolates showed inhibitory effects on *Salmonella typhi*, *E. Coli*, *Vibrio cholerae, Pseudomonas aeruginosa, Staphylococcus aureus*, and *Bacillus spizizenii*. Streptomycin was used as the positive control. The extracts had vigorous antibacterial activity, with *Streptomyces cyaneus* showing the most significant activity level. At 1.6 mg/mL, 0.8 mg/mL, and 0.4 mg/mL, inhibitory effects were seen. With a minimum inhibitory concentration (MIC) of 0.1 mg/mL, *Streptomyces cyaneus* had the lowest against *Vibrio cholerae*. The actinomycetes’ minimum inhibitory concentration (MIC) ranged from 0.8 mg/mL to 0.1 mg/mL.

Extracts’ impact on *S. typhi* development is depicted in Figure 5. After adding the extracts, the bacteria were cultured in Mueller Hinton Broth (MHB) in a microtiter plate. For the positive control, ciprofloxacin was used. By comparing the results from day 0 and day one following an 18-hour incubation period at 37°C, absorbance at 600 nm was obtained to create a growth curve. More investigation is required to determine the chemical structures in the extracts.

**Figure 5:**
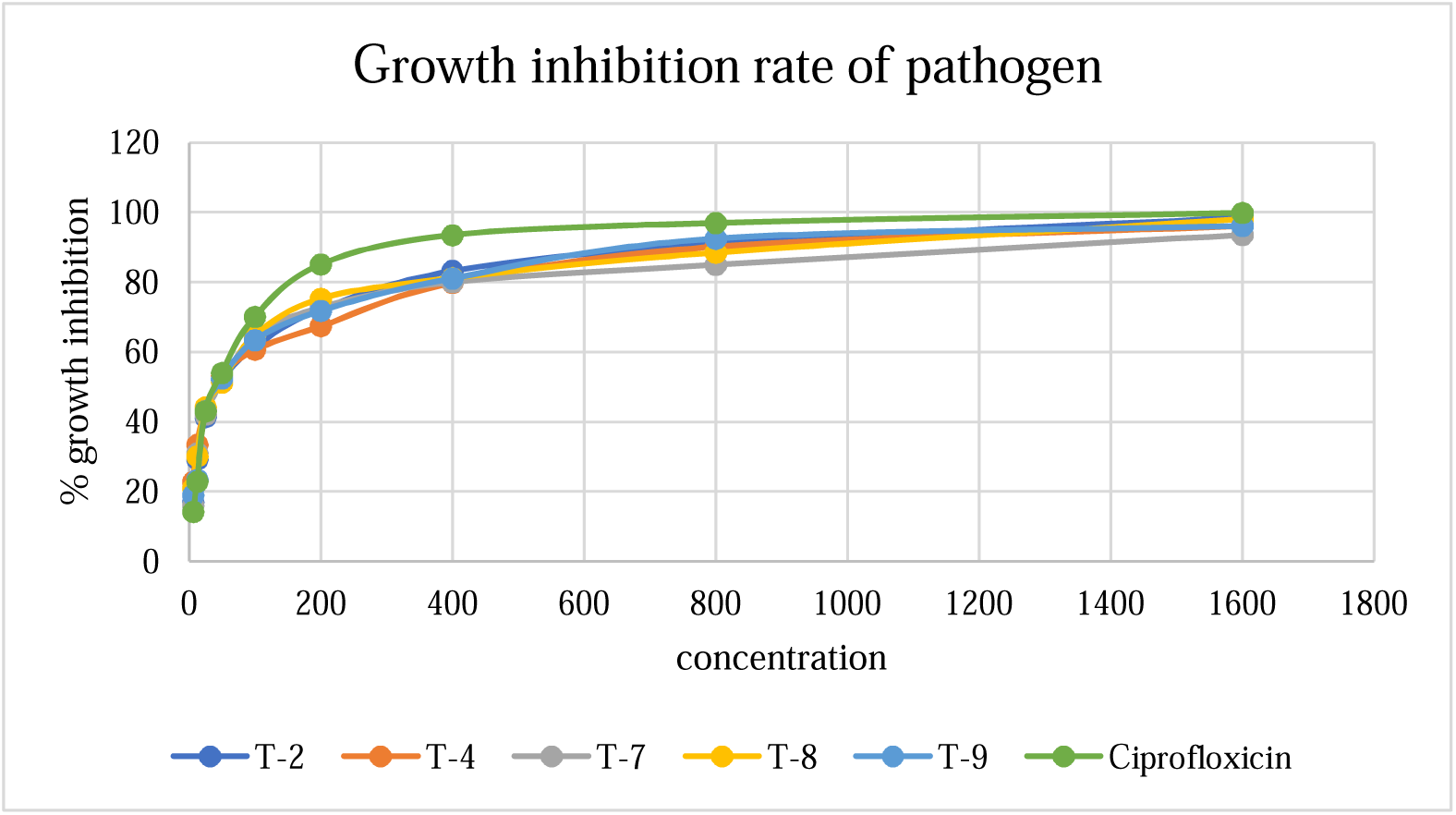
Determination of growth inhibition of the extracts against Salmonella typhi ATCC- 1408

## Antioxidant Activity

### The DPPH Free Radical Scavenging Assay

When antioxidant activity was assessed using DPPH free radical scavenging, all actinomycete extracts displayed concentration-dependent outcomes. Each extract exhibited antioxidant qualities; the highest IC_50_ value was recorded by *Streptomyces albogriseolus* (T-8) at 372.09±11.05 µg/mL, whereas ascorbic acid had an IC_50_ of 112.03±6.00 µg/mL. The IC_50_ values for each extract and standard are displayed in Figure 6, demonstrating that the extract’s scavenging activity is somewhat similar to the standard’s, suggesting modest antioxidant qualities.

**Figure 6:**
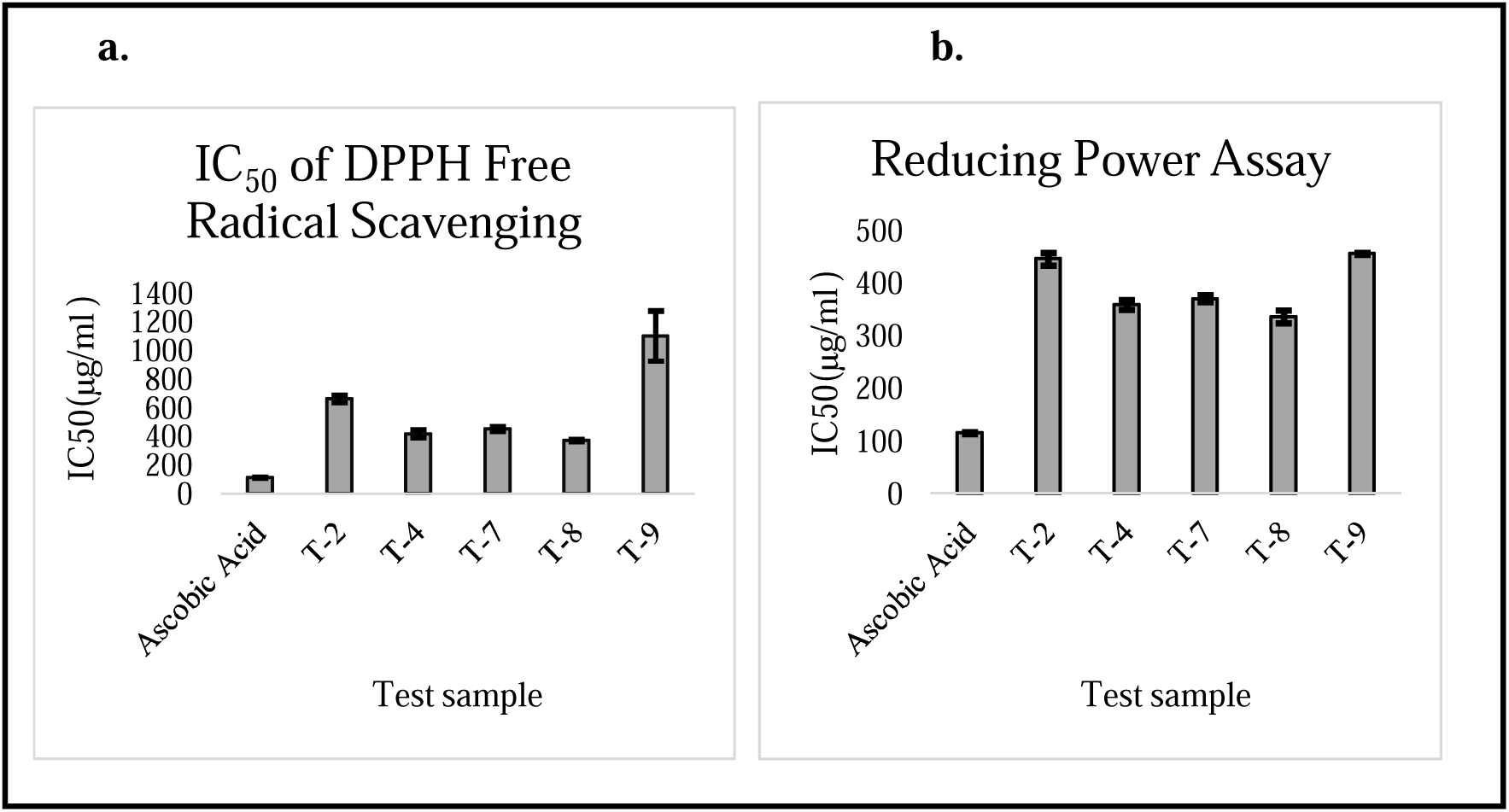
Antioxidant potential of actinomycetes extracts compared to the standard (ascorbic acid)

### Reducing Power Assay

Figure 6(b) presents the findings from the Reducing Power Assay for the standard (ascorbic acid) and extracts of actinomycetes.

The IC_50_ values for DPPH inhibition are shown in Figure 6(a) and are obtained by extrapolating the equation y = mx + c, where m is the slope and c is the intercept. The highest IC_50_ value was recorded by *Streptomyces albogriseolus* (T-8), at 372.09±11.05 µg/mL, while ascorbic acid had an IC_50_ of 112.03±6.00 µg/mL.

Using y = mx + c as a guide, IC_50_ values for power reduction are shown in Figure 6(b). Ascorbic acid’s IC_50_ was 114.97±2.99 µg/mL, but the extract from *Streptomyces albogriseolus* (T-8) exhibited the highest reducing power at 353.97±21.00 µg/mL. The reducing power experiment revealed that all extracts exhibited antioxidant characteristics by converting ferricyanide [Fe(CN)LJ]³LJ to ferrocyanide [Fe(CN)LJ]LJLJ in a concentration-dependent manner.

## Evaluation of Blood Coagulation Activity

### Serine Protease Inhibition Assay

The actinomycetes extract’ ability to inhibit serine protease was examined in this work; Figure 7 shows the IC_50_ values for each extract.

**Figure 7:**
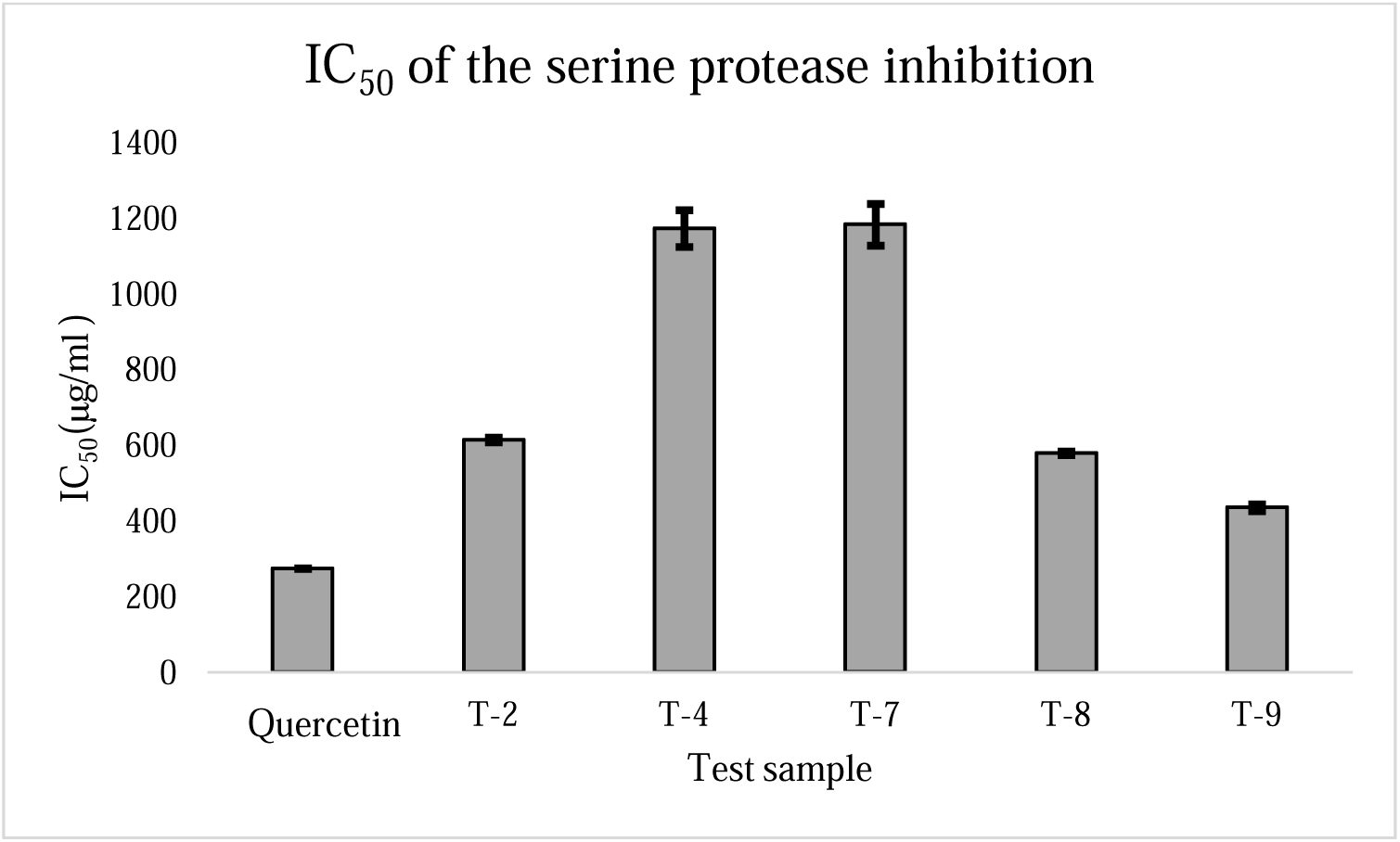
The IC_50_ values for the Serine Protease Inhibition assay are shown for both the actinomycete’s extracts and the standard quercetin.

Using the formula y = mx + c, the IC_50_ values for trypsin inhibition are displayed in Figure 7. The standard quercetin exhibited an IC_50_ of 274.54±1.26 µg/mL, whereas the T-9 extract had the highest value of 435.12±15.88 µg/mL, suggesting its efficiency among all extracts.

### Prothrombin Time Determination (PT) Assay

In the prothrombin time (PT) test, clotting time was affected by all extracts in a concentration-dependent manner. At 800 µg/mL, the standard Warfarin took the longest to clot- 40.26±1.11 minutes, while T-9 needed 12.08±1.46 minutes at the same dose. Clotting periods are given for blood, including standards and extracts at different concentrations, as shown in Supplemented Table 5.

## Anti-inflammatory Activity Assessment

### The Hypotonicity-induced Hemolysis Assay

The actinomycetes extract all performed similarly in the assay for hypotonicity-induced hemolysis. The IC_50_ values for the extracts and the standard (Aspirin) are shown in Figure 8. Both the extracts and the standard showed a concentration-dependent inhibitory effect.

**Figure 8:**
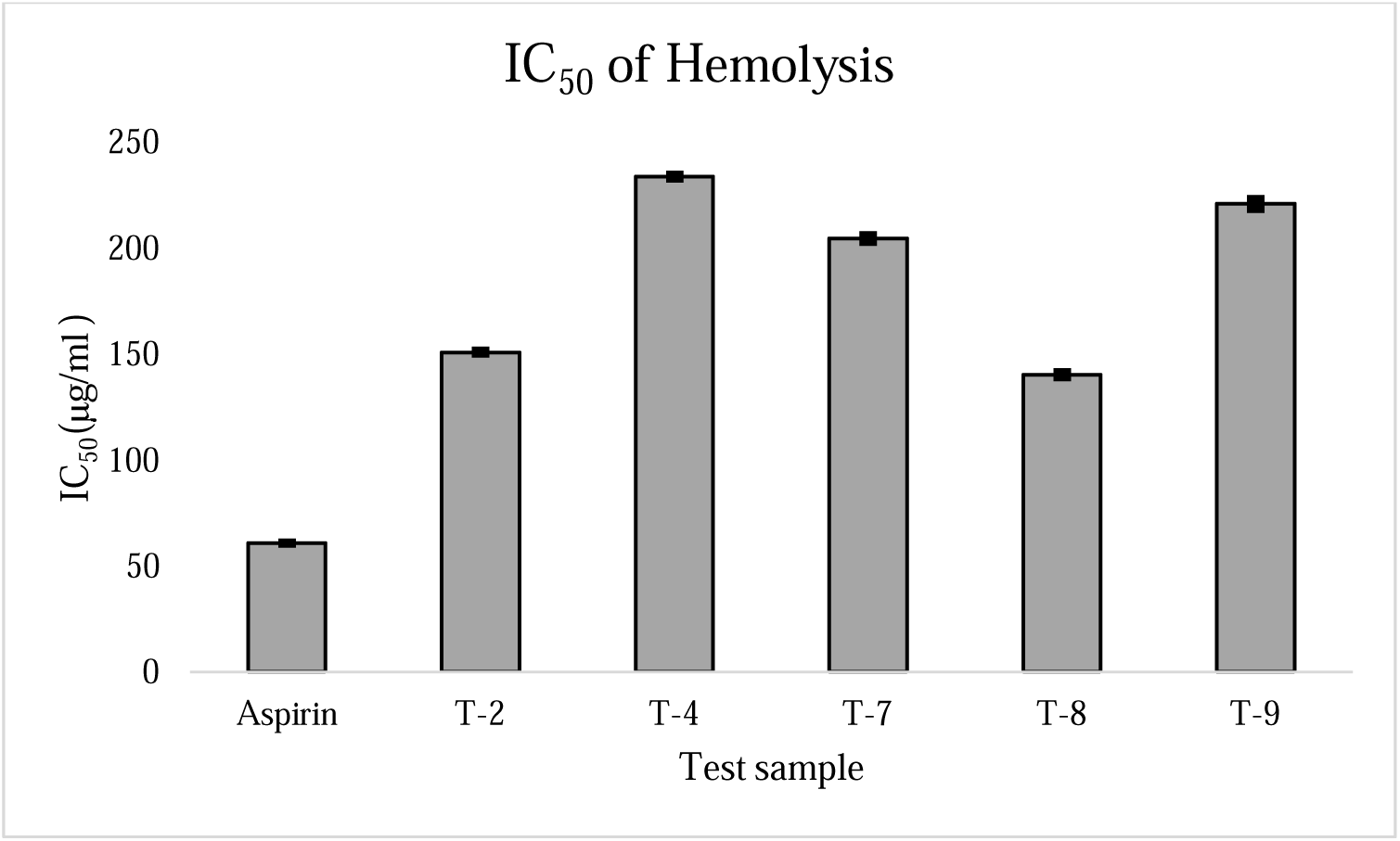
IC50 value of hypotonicity-induced hemolysis assay of actinomycetes extract and standard Aspirin in different concentrations.

Using the trendline equation y= mx+c, the IC_50_ value for hemolysis inhibition was ascertained. At 140.08 ± 2.30 µg/mL, the T-8 extract showed the most excellent IC_50_ value (Figure 8), whereas the IC_50_ value of the conventional Aspirin was 60.79 ± 0.76 µg/mL.

## Discussion

In this work, we examined Actinomycetes from sea sediments in Char Kukri Mukri and Char Montaz, Bangladesh’s Bhola district, using techniques that depended on culture. Actinomycetes strains 47 were isolated, 22 were chosen based on their morphology, and ten were set aside for additional examination. The majority of these isolates are Streptomyces, according to 16S rRNA gene sequencing.

Marine Actinomycetes are known for producing bioactive compounds (Duncan et al., 2015). Our research evaluated the crude secondary metabolites from various Actinobacteria strains for their antibacterial, anti-inflammatory, anticoagulant, and antioxidant properties. The antimicrobial tests demonstrated that extracts from five Streptomyces isolates showed activity. This finding supports the established ability of Streptomyces to generate diverse bioactive secondary metabolites. (Berdi, 2005; Bérdy, 2012).

This investigation’s techniques successfully separated Actinomycetes from maritime sediments. Comparably, Actinobacteria were identified as over 600 strains, primarily from the genera *Micromonospora, Kocuria, Nocardiopsis, Saccharomonospora,* and *Streptomyces*, after they were successfully isolated from maritime sediments in the Yellow Sea. Xiong et al. (2015)

In our study, bacterial isolates were initially identified based on morphological characteristics and then cultured in a selective Starch-Casein-Agar medium, pre-incubated at 30°C for 2 hours to ensure the viability of Actinomycetes. Subsequent 16S rRNA sequencing confirmed the species as follows: T-2 (*Streptomyces olivoverticillatus*), T-4 (*Streptomyces cyaneus*), T-5 (*Cytobacillus oceanisediminis*), T-6 (*Streptomyces albogriseolus*), T-7 (*Nocardiopsis synnemataformans*), T-8 (*Streptomyces albogriseolus*), T-9 (*Streptomyces atrovirens*), T-10 (*Cellulosimicrobium funkei*), T-11 (*Streptomyces labedae*), and A-2 (*Streptomyces camponoticapitis*). The sequences were submitted to GenBank.

With species confirmation, we fermented these isolates to produce secondary metabolites. The fermentation process was conducted in suitable media at 28°C for 48 hours under sterile conditions. After a four-week incubation period, we harvested a significant yield of secondary metabolites. Bacterial cells were separated via centrifugation, and the crude secondary metabolites were extracted using ethyl acetate (Undabarrena et al., 2016).

Bioactivity testing revealed that the extracts contained active metabolites, suggesting potential new biotechnological applications. The minimum inhibitory concentrations (MIC) of the extracts ranged from 0.8 mg/mL to 0.1 mg/mL, demonstrating significant antifungal activity, especially in extracts from *Streptomyces cyaneus*, consistent with Biswas and Solanki’s findings using ciprofloxacin as a positive control. Besides antibacterial properties, the extracts exhibited antioxidant, anti-inflammatory, and anticoagulant activities. Antioxidant activity was selectively evaluated through the Reducing Power Assay and DPPH Free Radical Scavenging Assay. One isolate, T-8, showed moderate antioxidant activity with IC_50_ values of 372.09±11.05 µg/mL and 353.97±21.00 µg/mL, respectively. Previous studies have also highlighted antioxidant-active secondary metabolites in Streptomyces strains from marine sediments (Janardhan et al., 2014; Padma B. et al., 2018).

The anticoagulant activity was confirmed through the Serine Protease Inhibition Assay, with an IC_50_ value of 435.12±15.88µg/mL for sample T-9, and the Prothrombin Time Determination Assay. Standard Warfarin inhibited blood coagulation for 40.26±1.11 minutes, while sample T-9 inhibited coagulation for 12.08±1.46 minutes, demonstrating its moderate anticoagulant activity, previously validated by 6 (Saniarahan & Sabah, 2006)

The anti-inflammatory activity of the five samples was evaluated using the Hypotonicity-Induced Hemolysis Assay. The sample T-8 exhibited the highest 50% inhibition (IC_50_) of hemolysis at 140.08±2.30 µg/mL, demonstrating moderate anti-inflammatory effects of the bacterial extract. Furthermore, studies have documented that *Actinomadura sp.,* isolated from marine sediments, produces metabolites with significant anticancer and anti-inflammatory activities.(Rangnekar & Khan, 2015; Selim et al., 2021)

Research explicitly contradicting the bioactivity claims of Actinomycetes secondary metabolites is uncommon. Baltz (2017) provided a broader perspective on challenges and limitations in the field without opposing the concept of secondary metabolite bioactivity.

Although the samples in this study were crude, many other studies have shown that the crude secondary metabolites of Actinomycetes possess antimicrobial and even anticancer activities (Ribeiro et al., 2020). In this study, not all isolates exhibited potent anticoagulant, antioxidant, or anti-inflammatory activities, but compounds with moderate activities were identified. Compounds with moderate anticoagulant activity can be beneficial in treating Peripheral Artery Disease (Gerhard-Herman et al., 2017). Secondary metabolites from Actinomycetes with moderate antioxidant activity are valuable in preserving skin care products and food (Fereidoon & Ying, 2010; Pullar et al., 2017). Furthermore, Actinomycetes’ secondary metabolites have a modest anti-inflammatory effect, suggesting that they may be able to treat neurodegenerative illnesses like Parkinson’s and Alzheimer’s. (Heneka et al., 2015).

The research, from sample collection to biochemical assays, was conducted with three replicates per assay to ensure the accuracy of the results. Cycloheximide and vancomycin were added to the Starch-Casein- Agar medium to prevent contamination. To maintain high assay precision, all equipment and reagents were obtained from reputable sources. Notably, the *Streptomyces cyaneus* extract displayed antimicrobial solid activity against *Vibrio cholerae*, highlighting its potential for antibiotic development.

Despite recent studies on biosurfactant production by microorganisms, marine environments have yet to be extensively explored. (Mukherjee et al., 2009) . This research could be more impactful if pure metabolites were isolated through partial purification. Before these compounds can be used as bioactive agents in living organisms, cytotoxicity testing is necessary. Optimizing the solvent could lead to medicinal formulations, and advanced bioinformatics techniques, such as docking and simulation, could be applied to these isolated compounds to enhance their potential applications.

## Conclusion

The coastal ecosystem is a largely untapped source of actinomycetes, which can be easily isolated from marine sediments and rhizosphere areas. These microorganisms offer a rich array of secondary metabolites. The results indicate that actinomycetes hold significant potential for discovering medically critical natural compounds. Specifically, actinomycetes from coastal sediments may serve as promising sources of antimicrobial agents. While this study analyzed a few activities, there are numerous other potential functions, such as anticancer, antidiabetic, cytotoxic, and hydrocarbon degradation, that warrant further investigation.

## Acknowledgement

We thank Dr. Md. Nazmul Hasan, the Director of the Genome Center at Jashore University of Science and Technology, for his invaluable assistance in sequencing the bacterial sample.

## Supplementary data

Additional data can be accessed online at FEMSMC.

